# Immune clustering reveals molecularly distinct subtypes of lung adenocarcinoma

**DOI:** 10.1101/2023.04.17.537140

**Authors:** Yan Lender, Ofer Givton, Ruth Bornshten, Meitar Azar, Roy Moscona, Yosef Yarden, Eitan Rubin

**Affiliations:** Shraga Segal Dept. of Microbiology, Immunology & Genetics, Ben-Gurion University in the Negev; The Shmunis School of Biomedicine and Cancer Research, Tel Aviv University; Dov Brayer 11, Ashkelon 7835417, Israel; Department of Immunology and Regenerative Biology, Weizmann Institute of Science

**Keywords:** Immunotherapy, refractive tumors, clustering, subtypes

## Abstract

Lung adenocarcinoma, the most prevalent type of non-small cell lung cancer, consists of two driver mutations in KRAS or EGFR. In general, these mutations are mutually exclusive, and biologically and clinically different. In this study, we attempted to find if we could separate lung adenocarcinoma tumors by their immune profile using an unsupervised machine learning method. By projecting RNA-seq data into inferred immune profiles and using unsupervised learning, we were able to divide the lung adenocarcinoma population into three subgroups, one of which appeared to contain mostly EGFR patients. We argue that EGFR mutations in each subgroup are different immunologically which implies a distinct tumor microenvironment and might relate to the relatively high resistance of EGFR-positive tumors to immune checkpoint inhibitors. However, we could not make the same claim about KRAS mutations.

**Simple Summary:** Lung adenocarcinoma, the most prevalent type of non-small cell lung cancer, is most commonly driven by mutations in KRAS or EGFR. In this study, we attempted to find if we could separate lung adenocarcinoma tumors by their immune profile using an unsupervised machine learning method. We used established tools to infer the immune profile of each tumor from its RNA-seq and using unsupervised learning, we were able to divide the lung adenocarcinoma population into three subgroups, one of which appeared to contain mostly patients with EGFR mutations. We argue that tumors with EGFR mutations in each subgroup are different immunologically which implies a distinct tumor microenvironment and might relate to the relatively high resistance of EGFR-positive tumors to immune checkpoint inhibitors. However, we could not make the same claim about KRAS mutations.

## 1. Introduction

Lung cancer is the most common cause of death from cancer worldwide with 1.79 million deaths in 2020 [1]. Non-small cell lung cancer (NSCLC) accounts for 85% of all lung cancers, with 40% of the cases being lung adenocarcinoma (LUAD), 25–30% squamous cell carcinoma, and 5–10% large cell carcinoma [2]. The five-year survival rate of LUAD patients is about 15%, lower than the overall NSCLC five-year survival rate (18%), as it is usually diagnosed in a metastatic form [3].

The most common driver mutations in LUAD are KRAS and EGFR (∼30%, and ∼15%, respectively in western populations) [4,5]. The most common mutation occurs in KRAS, a member of the Rat sarcoma (RAS) family. It most commonly involves a gain-of-function mutation (usually resulting from a single base substitution) that activates KRAS, promoting cancer invasion and metastasis. Patients harboring a KRAS mutation usually respond poorly to chemotherapy but are more likely to respond well to immunotherapy [6,7] than patients with the EGFR mutation. It was shown in mice that KRAS mutations are related to the increased CD8+ T cell, regulatory T cell, and myeloid cell migration into the tumor [7]. In addition, KRAS-mutated patients carry, on average, a higher tumor mutation burden than EGFR-mutated patients [8].

EGFR is the second most common driver mutation in LUAD. This mutation encodes a transmembrane glycoprotein that plays a major role in the cell decision between proliferation and apoptosis [9,10]. The two most common mutations found in the EGFR gene in LUAD tumors include exon 19 in-frame deletions and leucine-to-arginine substitution in exon 21 [10-12]. Together, these two mutations account for 85–90% of all EGFR mutations [11-15] and are associated with low mutation burden [12].

From a clinical perspective, EGFR mutations usually are common in non-smoking females of Asian ethnicity [13], while KRAS mutations are more common in smokers [14], but no differences were detected in the prognosis and diagnosis of the two mutations [15] Since both mutations are clinically too similar upon diagnosis, we sought to find a way to distinguish between them by examining the immune repertoire of patients carrying these mutations. We constructed a pipeline by using unsupervised learning based on “xCell” (see methods below) [16]. We found that LUAD patients from The Cancer Genome Atlas (TCGA) could be divided into three distinct groups. To test if the groups were molecularly distinct, we checked if driver mutations were randomly distributed across the groups. Most EGFR-mutated patients fell into one cluster, suggesting a possible way to characterize LUAD groups using unsupervised learning.

## 2. Materials and Methods

### Study population

This study was conducted on The Cancer Genome Atlas population, which has been described elsewhere [17]. A brief description of the study population is provided in Table 1 below. Mutation data were available for 566 patients and clinical data for 498 patients.

**Table 1.** Demographic characteristics of the TCGA LUAD populations

| Characteristics |  | EGFR status |  | P-val |
| --- | --- | --- | --- | --- |
|  |  | WT | Mut | WT-Mut |
|  | **Population (n)** | 492 | 66 |  |
| **Tumor grade** | Grade I | 247 | 29 | 0.65 |
|  | Grade II | 105 | 14 | 0.89 |
|  | Grade III | 70 | 13 | 0.42 |
|  | Grade IV | 21 | 6 | 0.19 |
|  | Not available | 49 | 4 |  |
| **Smoking** | Yes | 386 | 34 | 0.07 |
|  | No | 46 | 27 | 3.26E-08 |
|  | Not available | 60 | 5 |  |
| **Diagnosis age (years)** | <=60 | 140 | 17 | 0.84 |
|  | >60 | 289 | 42 | 0.78 |
|  | Not available | 63 | 7 |  |
| **Gender** | Female | 230 | 43 | 0.14 |
|  | Male | 215 | 19 | 0.16 |
|  | Not available | 47 | 4 |  |
| **Race** | White | 336 | 46 | 0.99 |
|  | Black | 46 | 6 | 0.87 |
|  | Asian | 5 | 3 | 0.1 |
|  | Native American/Native Alaskan | 1 | 0 | 1 |
|  | Not available | 104 | 11 |  |

### Data

LUAD mutation data were downloaded from cBioPortal [18,19] on 2.2.2021. Before analysis, outliers were removed based on their mutation count, following z-score normalization (z>3.5) based on the mutation count we have chosen a little higher value than the default used (3) [20]. The Z-score was calculated for LUAD EGFR- and KRAS-mutated patients using single nucleotide variation (SNV) data [21] for total mutation calculation, downloaded on 20.7.2020. Seven samples ‘05-4382’, ‘17-Z022’, ‘55-7907’, ‘55-8506’, ‘55-A490’, ‘78-7155’, and ‘86-A4JF’ were removed for having exceptionally high Z-scores. All subsequent analyses were performed after this removal.

For k-means clustering, the number of clusters was optimized by the knee method [22]. The random seed was set to 42 during the calculations.

For clinical information, data was downloaded on 23.5.21 from cBioPortal [18,19] In addition, EGFR patients’ smoking information was downloaded from Xena GDC [23] (27.5.2021).

Overall survival information was downloaded from cBioPortal [23] on 23.5.21.

### Statistics

To test for differences in clinical properties between the clusters, a chi-square contingency test (Fisher’s exact test) was used to test for the independence of each property in the EGFR wild-type (WT) groups. Testing was performed using the Python programming language (ver. 3.8), with the SciPy library (version 1.7.3). For differential expression analysis, the R programming language (version 4.2.0 [24]) was used with the Deseq2 package (version 1.36.0 [25])

### xCell

To infer immune repertoire from bulk mRNA profiles, we used xCell, a computational method that estimates the cell-type abundance of immune and stromal cell types from transcriptomic data. The method is based on an algorithm that compares the expression levels of signature genes that are specific to each cell type with the expression levels of all other genes in the dataset.

### Immune Inference from Bulk Expression Profiles

To infer immune infiltration from RNA-seq data expression profiles, we used TIMER (ver. 2.0) [26-32], which uses cell-type specific gene signatures using the default parameters (downloaded in bulk on 13.4.22, LUAD extracted). The normalized RNA-seq profile (see Methods above) was converted to immune-related cell type content using xCell. Four EGFR patients lacked expression data and were removed. The data were scaled using Z-score transformation and clustered using the K-means algorithm. Both steps were performed with the scikit-learn library (version 1.1.1) [33] in Python. Only clusters with five or more individuals were maintained. K-means clustering was used to divide the patients, and the resulting clusters were compared (Figure 1). The same pre-processing and clustering procedure was performed on raw expression data of LUAD (downloaded from FirebrowseR [34]. on 13.4.2022), without immune conversion.

**Figure 1.**
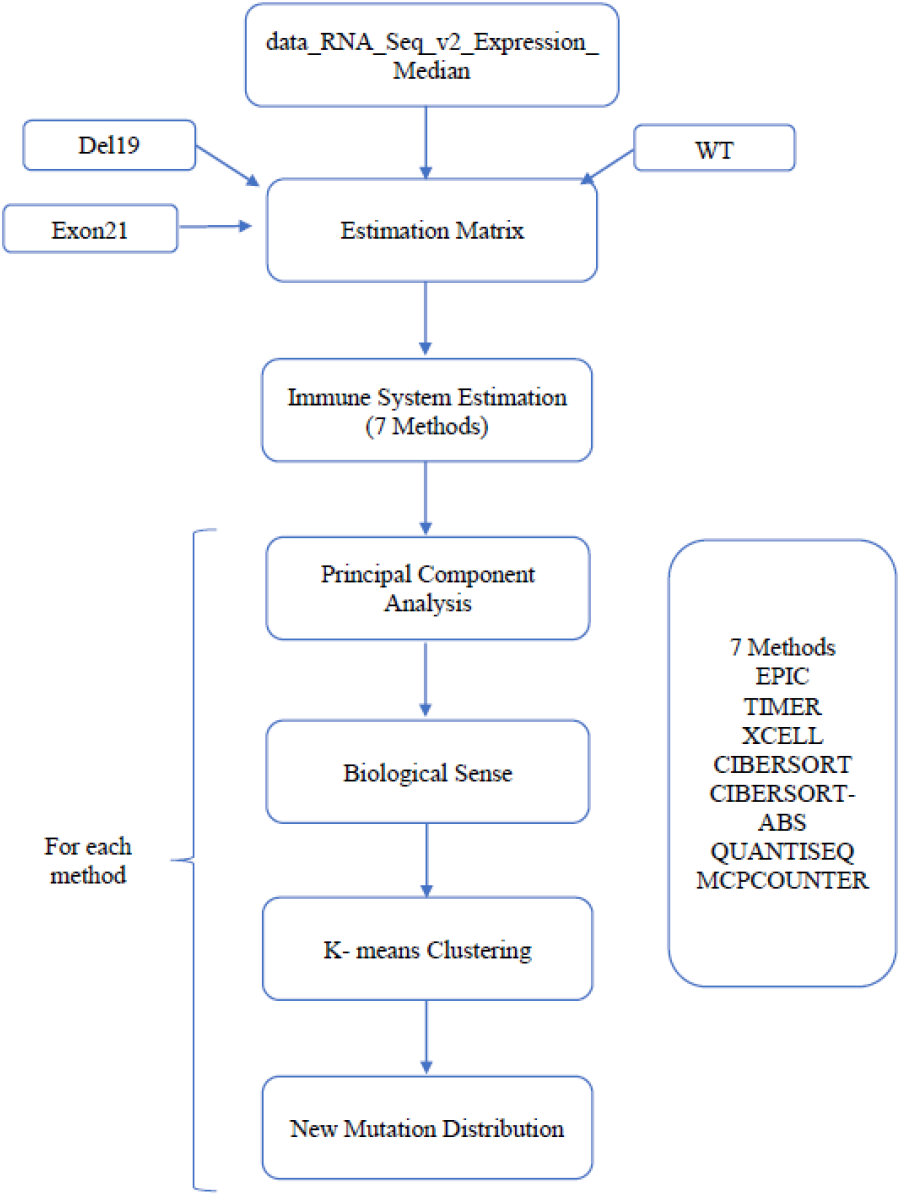
The way from expression profiles to new mutation distribution (based on previous knowledge). This figure describes the pipeline utilized in this work to examine the value of immune inference to find new ways to divide LUAD patients. The new mutation distribution was calculated for each cluster separately.

### Overall survival

Kaplan Meier curve has been drawn using Matplotlib package [35] in Python. Out of 559 patients, 61 were omitted for missing overall survival data (n=498). A pairwise log-rank test has been applied to find differences between the populations.

### Differential expression analysis

The expression of EGFR-mutated patients was compared between patients assigned to sub0 (n=33), sub1 (n=4), and sub2 (n=25) using DeSEQ2. In addition, differential expression (DE) was performed on mutated EGFR between nsubx to nsubx-4 where nsubx is sub2 or sub0 patients after removing four random patients.

### Availability

All the codes and data described in this work are available upon request.

## 3. Results

### Characteristics of the Study population

This study was based LUAD subsection of TCGA. The population used in this data set is described in detail elsewhere [27] (see also Table 1 for a description of the population characteristics). Briefly, it includes 558 patients (after integration with mutation data), most of whom were diagnosed at age of >60 (n=289). Dividing this population based on EGFR status revealed little gender difference in the EGFR-WT subpopulation (n=230 vs. 215). In contrast, females are significantly over-represented in EGFR-mutated patients compared to EGFR-WT patients (n=43 vs. 19; p=0.02, chi-square test, data not shown), as previously reported [28]. In terms of ethnicity, this study focused on Caucasians (n=336), although ethnicity is unknown for ∼25% of the population. The diagnosis was mostly at first grade (n=247), but a weak non-significant negative correlation is observed between the stage and frequency of detection (corr = -0.63, p=0.09, Spearman). There are significantly more non-smokers in the EGFR-mutated group than in the WT group (n=27/66, 46/492, respectively, p-value<0.05, chi-square test), in accordance with previous reports [28]. Overall survival was not significantly different in the WT group than in the EGFR-mutated group (p=0.1, log-rank pairwise test) (Table 1).

### Immune inference and unsupervised learning detect new groups

To infer the immune status in the tumor from bulk mRNA profiling, we used the xCell tool for this purpose. Multiple methods have been considered to perform this task, and the choice of xCell was mostly arbitrary.

First, we tested the hypothesis that using inferred immune status would reveal groups missed when raw expression profiles are used. For this, we used the K-means clustering algorithm [29], with the original expression profiles (based on RNA-seq) and the knee analysis method to decide on the number of groups [22]. The population was divided into four subgroups: sub0, sub1, sub2, and sub3. KRAS mutated patients were non-randomly distributed among these groups (p=0.01, chi-square test), with sub1 enriched with KRAS-WT patients. On the other hand, no enrichment in EGFR-mutated patients could be detected (p=0.18, chi-square test).

These results were compared to clustering with the same methodology (K-means clustering with K=3, knee method) using inferred immune cell content instead of raw expression. K-means assigned patients to three subgroups: sub0, sub1, and sub2. As can be seen in Table 3, sub0 was highly and significantly enriched with EGFR-mutated patients: 33 patients in sub0 (20.1%) were EGFR-mutated, compared to 5.3% in sub1 and 9.5% in sub2 (p=6*10-4, chi-square test).

**Table 3:** The distribution of patients (by EGFR and KRAS status) in different patient groups. The distribution of mutations (EGFR and KRAS status) in different patient clusters. EGFR and KRAS mutations were counted after K-means clustering into “sub” groups. In some cells, the fraction of patients carrying a mutation is shown in parentheses.

|  | sub0 | sub1 | sub2 |
| --- | --- | --- | --- |
| EGFR-WT | 131 | 71 | 239 |
| EGFR-mutated | 33 (20.1%) | 4 (5.3%) | 25 (9.5%) |
| KRAS-WT | 178 | 59 | 116 |
| KRAS-mutated | 86 (32.6%) | 16 (21.3%) | 48 (29%) |

To assess whether these groups are clinically different, the survival of patients from the three groups was compared using the Kaplan–Meier plot (Figure 3). The groups showed distinct survival patterns: patients from sub2, which was enriched with EGFR-WT (n=260), had a significantly worse prognosis than those in the sub0 group (p-value=0.001, log-rank test) or in the sub1 group (9 patients were omitted due to lack of clinical data,p-value=0.04, log-rank test) (Figure 4)

### Differential expression of immune checkpoints between immune cell clusters

We next compared the expression of known immune checkpoints between the different groups. Since these groups were defined by the abundance of immune cells, it is possible they would also differ in immune checkpoint expression. Since these are the target of immune checkpoint inhibitors, we were interested in comparing their expression between the subgroups (data not shown). Significant differences were observed for four targets: TIGIT expression differed in all three comparisons: between sub1 and sub0 (p=0.01), between sub2 and sub0 (p=0.01), and between sub2 and sub1 (p=7*10-4). PD-L1 (CD274) expression differed in two comparisons: sub1/sub0 (p=2.54*10-7) and sub2/sub0 (p=0.004), as did CTLA4 (p=0.02 and p=0.003, respectively). PD-1 (PDCD-1) expression differed only between sub2 and sub1 (p=0.03). No other known immune checkpoints were significantly different between any of the clusters.

Finally, we tested the hypothesis that the four EGFR-mutated patients were wrongly assigned to sub1. If this were the case, we would expect little difference between the EGFR-mutated patients assigned to sub1 (n=4) and EGFR-mutated patients in other clusters. A differential expression (DE) comparison between the EGFR-mutated patients in each sub-group revealed significant results (p<0.05, FDR<0.25) in both comparisons: 1,428 genes significantly differed between EGFR-mutated patients from sub0 and EGFR-mutated patients from sub1, and 510 genes differed between sub2 to sub1 patients. For comparison, four patients were randomly chosen from sub0 or sub2 and compared, using the same procedure, to the remainder of the EGFR-mutated patients in the same cluster (see Methods). This process was repeated 18 times, to give the average and standard deviation of the differences expected if these four patients in sub1 had mistakenly been assigned to it. Fewer genes were significantly different within either of the subgroups (77±149.2 and 117±266.8 results were obtained for sub0 and sub2, respectively), suggesting that EGFR-mutated patients were randomly assigned to sub1, although the difference was significant for sub0 (p<1.23*10-18, z-test), but not for sub2 patients (p=0.15, z-test).

## 4. Discussion

In this work, we used inferred immune content rather than expression patterns to divide LUAD patients. This approach revealed clusters that are more strongly associated with mutation type (Table 3), and that are distinct in terms of survival (Figure 4), differentially expressing immune checkpoint genes (Figure 5).

The immune uniqueness of tumors harboring EGFR or KRAS mutations in terms of immune response is poorly characterized, but the differences in the demographics of the two populations have been reported [13,14]. When it comes to treatment, LUAD patients are divided by their driver mutations although the therapeutic usefulness of the traditional division in the face of immunotherapy has recently been challenged [31]. The use of immunotherapy dramatically increases the need for a new and more precise way to divide LUAD patients. To the best of our knowledge, this is the first time an unsupervised machine-learning method has been applied to divide LUAD patients in TCGA based on their inferred immune profiles.

Since the clustering approach presented here is based on (inferred) immune profiles, it is not surprising to see large differences in the expression of immune checkpoints. Accordingly, PDCD1 (also known as PD-1), which is expressed in activated T cells, was overexpressed in sub1 compared to sub0. This may indicate that there were more activated T cells in the tumors assigned to this cluster. Accordingly, inhibiting the PD-1/PD-L1 axis, the main target of most immune checkpoint inhibitors that are currently used, is expected to be more helpful to patients in the sub1 cluster. This is in agreement with our finding that sub1 tumors rarely carry EGFR mutations. Immune checkpoint inhibitors were shown to be less valuable for patients with EGFR mutations [31]. In other words, tumors with EGFR mutations do not attract the immune response that EGFR-WT tumors attract, and as a result, they are both less susceptible to immune checkpoint inhibitors (ICIs) and tend to fall into unique immune profile clusters. EGFR-mutated tumors were grouped mostly in sub0, and they constituted more than 20% of that group (Table 3). The finding that some of the EGFR-WT tumors clustered together with EGFR-mutated cases suggests that they attract the same immune response as EGFR-mutated tumors. This is in according that most EGFR-WT patients do not respond to ICIs [32]. Those EGFR-WT tumors that do not respond to ICIs may be more similar in their immune profile to EGFR-mutated patients, which too often are restrictive to ICI treatment. The division we present here may be the first step toward finding a division that can guide ICI therapy based on an immune profiling approach.

Our study population, taken from TCGA, is consistent in some aspects with previous reports. For example, EGFR mutations tend to occur more frequently in patients who never smoked than in smokers [36]. Unexpected differences were found, as previously reported, in the overall survival of both the KRAS- and EGFR-mutated groups versus the WT group (Figures 1 and 2). These differences are proposed to result from a biased LUAD TCGA population in which most of the patients had the disease identified at an early stage. However, in other aspects, the TCGA population is abnormal. The overwhelming majority of patients included in this study were diagnosed in Stage 1, while most lung adenocarcinomas are usually diagnosed at later stages [37].

**Figure 2.**
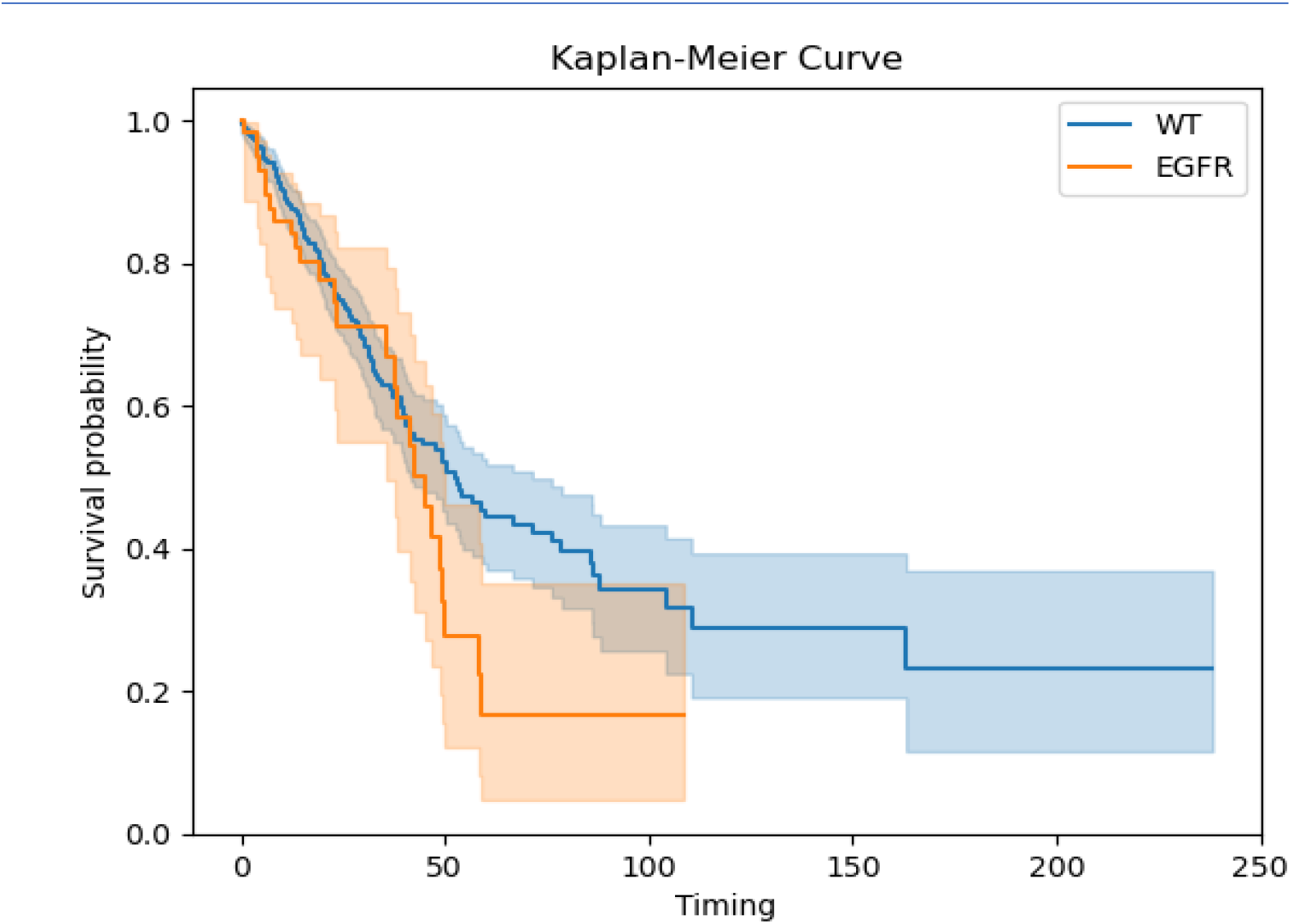
Kaplan-Meier curve of EGFR-mutated and EGFR-WT patients. The Y-axis shows the survival probability, while the X-axis represents the respective time interval. Separate lines, in different colors, each with a band of the same color, represent the comparison of EGFR-WT (n=437) vs. EGFR-mutated patients (n=61), *p*=0.1; logrank test.

**Figure 3.**
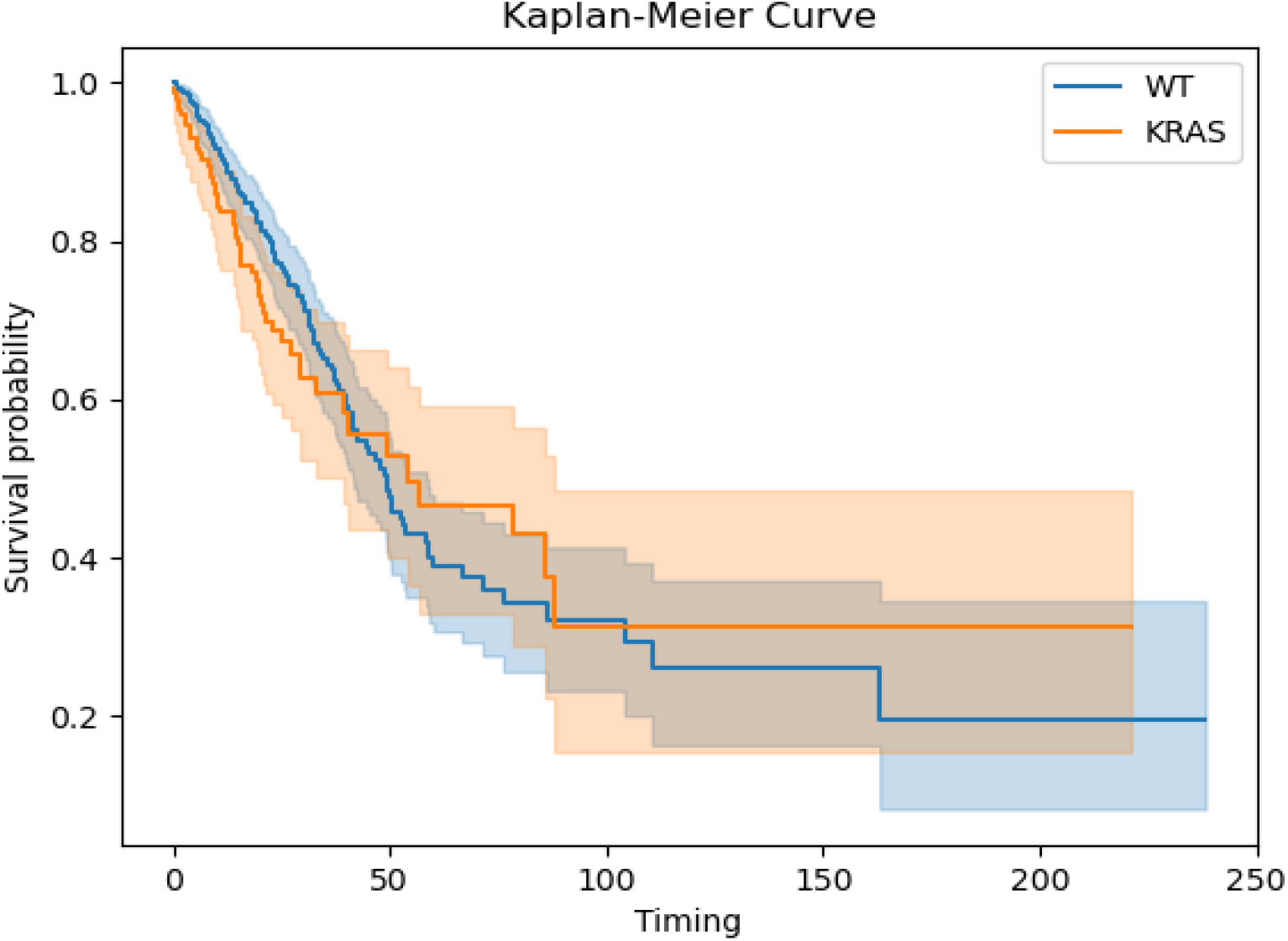
Kaplan-Meier curve of KRAS-mutated and KRAS-WT patients. The Y-axis shows the survival probability, while the X-axis represents the respective time interval in months. Separate lines, in different colors, each with a band of the same color, represent the comparison of KRAS-WT (n=347) vs. KRAS-mutated patients (n=151), p=0.45 log-rank test.

**Figure 4.**
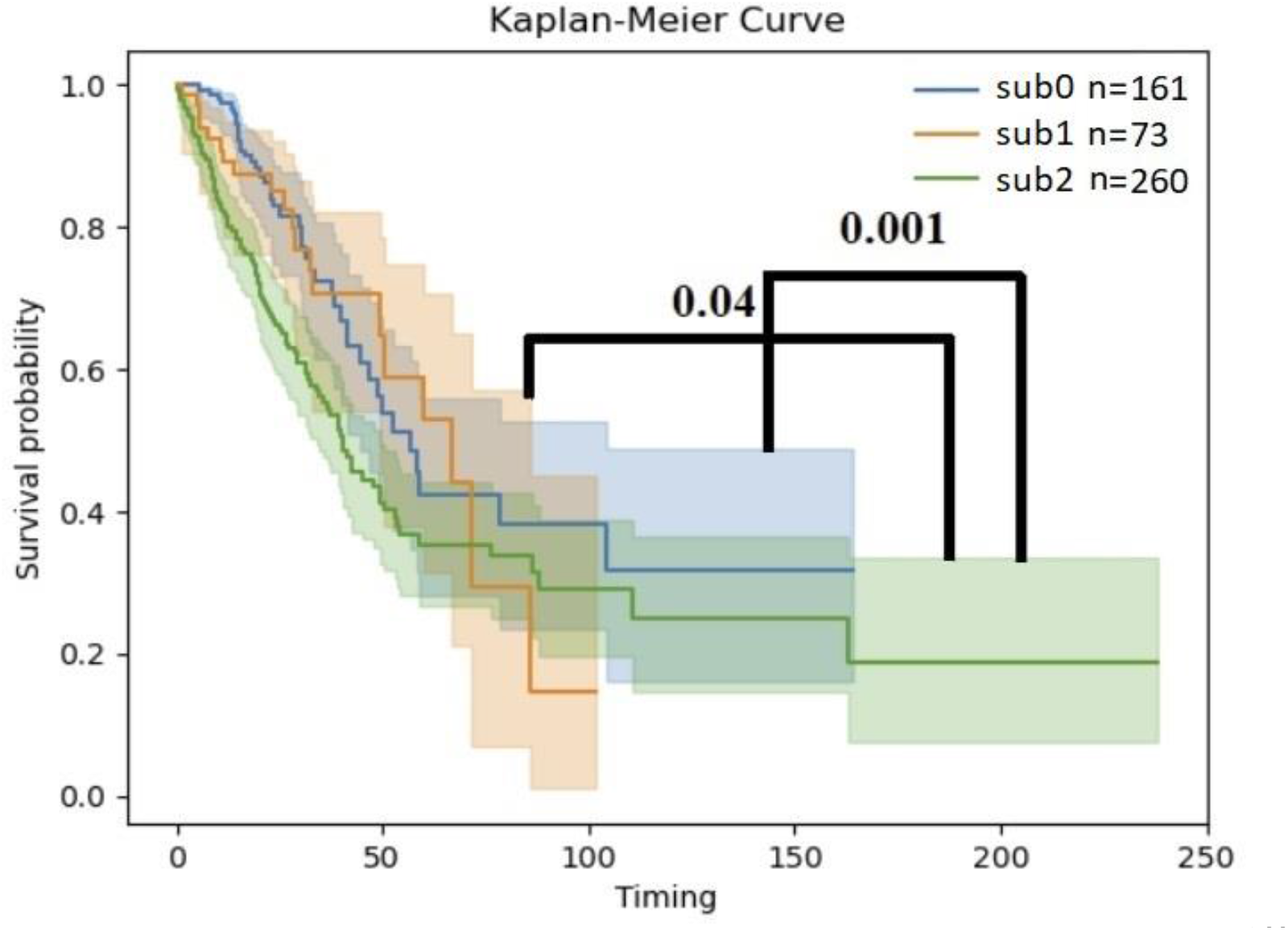
Kaplan-Meier curve of sub0, sub1, and sub2 groups of patients. Statistical comparisons, performed with the log-rank method [30] are shown as black lines. Significant differences were found for sub0 vs. sub2 (*p*=0.001, log-rank test) and sub1 vs. sub2 (*p*=0.04, log-rank test), but not for sub0 vs. sub1 (*p*=0.87, log-rank test).

**Figure 5.**
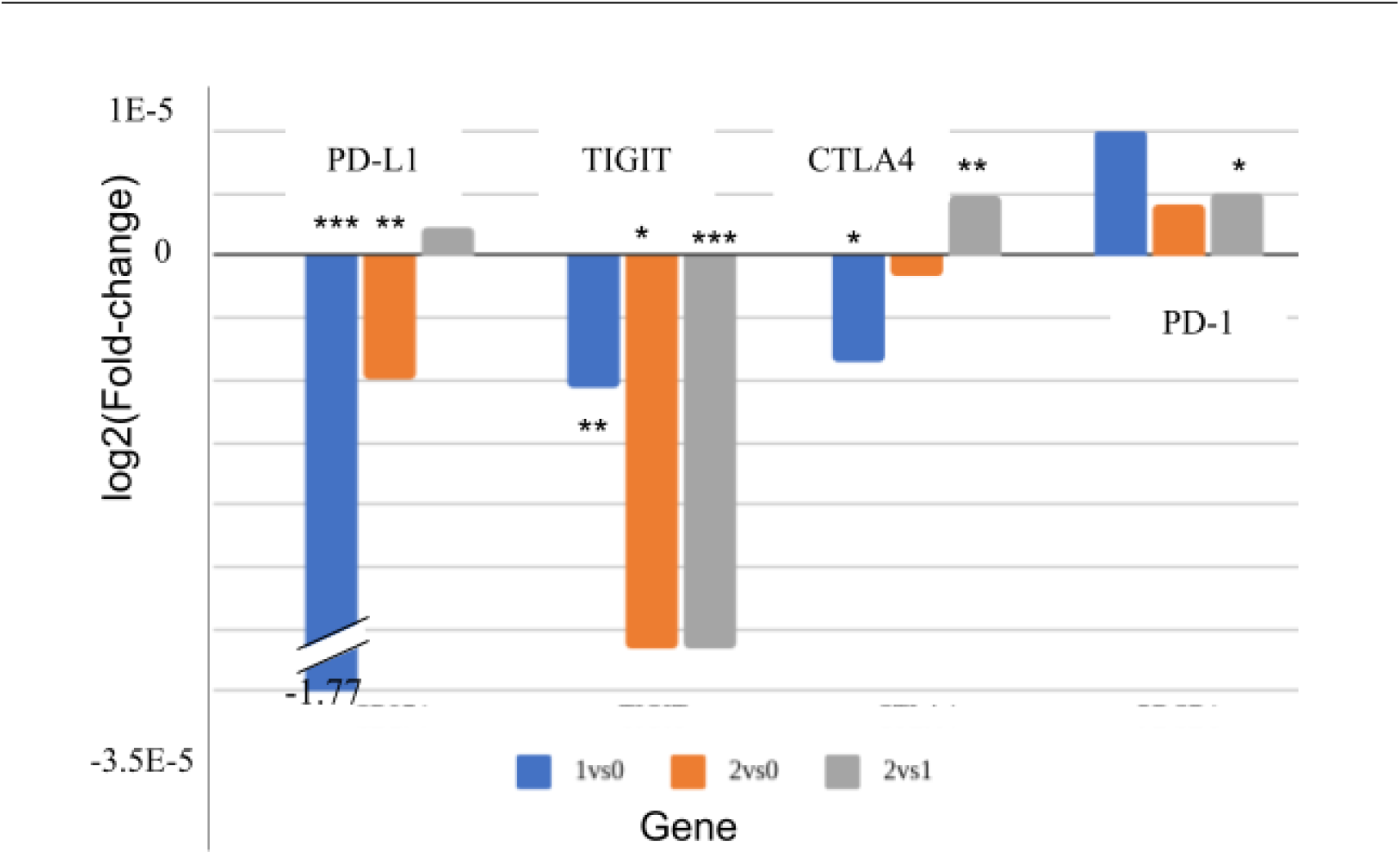
Differences in immune checkpoint expression between immune-defined subgroups. The values are minus log normalized with base 2 of the mean fold change. Asterisks indicate the significance of the difference (***p<0.001; **p<0.01; *p<0.05).

This work is based on unsupervised machine learning (i.e. clustering) [38,39]. Unlike previous works, we used inferred immune cell estimates to cluster tumors together. We used xCell to project RNA-seq-based expression profiles into inferred immune profiles. While numerous other methods have been proposed to infer immune cell content from bulk RNA-seq [40–42], a complete description of all these methods is beyond the scope of this work. However, repeating the analysis with other immune inference methods gave similar results (see Table S1), suggesting that our findings do not depend on the choice of the immune inference method.

We bring several lines of evidence that support the conclusion that classifying patients by inferred immune profiles divides them into biologically meaningful groups: (i) When patients are clustered based on expression without immune inference, a weak non-random bias in KRAS-positive patients to one particular cluster is observed (Table S2), while EGFR-positive patients were randomly distributed among the clusters. On the other hand, when inferred immune cell counts were used for clustering, EGFR-mutated patients, but not KRAS-mutated patients, were distributed non-randomly between the clusters with a more distinct bias (Table 3: sub2 vs. sub0, p=0.002; sub1 vs. sub0, p = 0.003). (ii) The clusters formed with immune inference differed in overall survival (Figure 4), with sub2 differing significantly from sub1 and sub0. (iii) Highly significant differences were observed in the expression levels of some of the targets of immune checkpoint inhibitors (e.g. PD-L1, which differed between sub1 and sub0) (Figure 5). It should be noted that both EGFR and KRAS mutation information has been taken from LUAD TCGA, and they are mutually exclusive, hence why the population number is the same for both groups.

Based on previous reports [43], enhanced PD-L1 expression on tumor cells leads to T-cell exhaustion, promoting tumor proliferation and survival [44]. Thus, an immune check-point inhibitor (e.g. anti PD-L1) may assist in releasing the T cells from exhaustion. High PD-L1 expression has been approved by The Food and Drug Administration (FDA) as a biomarker [45]. We propose that the patients from this sub0 cluster are less likely to respond to ICIs that target the PD1/PD-L1 axis, but the other clusters may respond to anti PD1/PD-L1 inhibitors.

This study is largely based on the accuracy of immune inference of xCell as a classification. Other immune inference methods have not been rigorously pursued.

Methods for directly measuring immune profiles have been suggested [46,47]. It would be interesting to examine clustering on actual, rather than inferred, immune profiles to cluster LUAD patients.

## 5. Conclusions

To conclude, we show that immune inference can uncover new clusters of LUAD patients that are otherwise hidden. We directly show that using immune inference gives more compact subgroups and that this division is clinically meaningful. We also show that these clusters differ molecularly, at the very least, in the expression of ICI targets. Specifically, we show that PD-L1 is markedly lower in one cluster while TIGIT is lower in another.

## Supplementary Materials

The following supporting information can be downloaded at: www.mdpi.com/xxx/s1, Table S1: PCA of other methods; Table S2: Comparison of clustering with and without immune inference.

## Author Contributions

Conceptualization, Y.L., Y.Y and E.R.; methodology, Y.L., E.R., O.G., R.B and R.M.; software, M.A.; investigation, Y.L., Y.Y. and E.R.; writing—original draft preparation, Y.L and E.R..; writing—review and editing, Y.L, E.R. and Y.Y; supervision, E.R. and Y.Y.; funding acquisition, E.R. All authors have read and agreed to the published version of the manuscript.

## Data Availability Statement

All the data available upon reasonable request

## Acknowledgments

The results published here are in whole or part based upon data generated by the TCGA Research Network: https://www.cancer.gov/tcga

## Conflicts of Interest

All authors have nothing to declare.

**Table S1:** PCA of other methods. Because each method uses its own way to estimate the immune profile, the score may be arbitrary. Positive scores indicate an immune impact. The higher the score, the higher the impact and vice versa.

| Cell type | EPIC | TIMER | CIBERSORT | CIBERSORT-ABS | QUANTISEQ | MCPCOUNTER | XCELL |
| --- | --- | --- | --- | --- | --- | --- | --- |
| **PC1** |  |  |  |  |  |  |  |
| Cancer associated fibroblast | 0.59 | ND | ND | ND | ND | 0.14 | ND |
| T cell CD8+ | -0.49 | 0.31 | -0.25 | 0.36 | -0.23 | 0.36 | 0.24 |
| T cell CD4+ | -0.34 | 0.47 | ND | ND | ND | ND | ND |
| NK cell | 0.05 | ND | ND | ND | -0.16 | ND | ND |
| Endothelial cell | 0.05 | ND | ND | ND | ND | 0.12 | ND |
| B cell | 0 | 0.48 | ND | ND | -0.24 | 0.27 | 0.25 |
| Macrophage | 0 | 0.14 | ND | ND | ND | ND | ND |
| Neutrophil | ND | 0.34 | ND | ND | -0.17 | -0.11 | ND |
| Mast cell activated | ND | ND | 0.42 | ND | ND | ND | ND |
| Myeloid dendritic cell resting | ND | ND | 0.35 | ND | ND | ND | ND |
| Monocyte | ND | ND | 0.34 | ND | 0.09 | 0.47 | ND |
| Macrophage M2 | ND | ND | 0.3 | 0.31 | -0.31 | ND | ND |
| NK cell resting | ND | ND | -0.27 | 0.31 | ND | ND | ND |
| Macrophage M1 | ND | ND | -0.24 | 0.39 | -0.4 | ND | ND |
| Macrophage M0 | ND | ND | -0.22 | ND | ND | ND | ND |
| T cell follicular helper | ND | ND | -0.22 | ND | ND | ND | ND |
| T cell CD4+ memory activated | ND | ND | -0.21 | 0.23 | ND | ND | ND |
| T cell CD4+ memory resting | ND | ND | ND | 0.34 | ND | ND | ND |
| T cell regulatory (Tregs) | ND | ND | ND | 0.28 | -0.46 | ND | ND |
| B cell memory | ND | ND | ND | 0.24 | ND | ND | 0.22 |
| B cell naive | ND | ND | ND | 0.21 | ND | ND | ND |
| Myeloid dendritic cell | ND | ND | ND | ND | 0.23 | 0.25 | 0.24 |
| Macrophage/Monocyte | ND | ND | ND | ND | ND | 0.47 | ND |
| T cell CD8+ central memory | ND | ND | ND | ND | ND | ND | 0.26 |
| T cell CD4+ naive | ND | ND | ND | ND | ND | ND | 0.23 |
| Plasmacytoid dendritic cell | ND | ND | ND | ND | ND | ND | 0.23 |
| Class-switched memory B cell | ND | ND | ND | ND | ND | ND | 0.21 |
| **PC2** |  |  |  |  |  |  |  |
| Endothelial cell | 0.53 | ND | ND | ND | ND | -0.31 | ND |
| B cell | 0.46 | -0.38 | ND | ND | -0.52 | 0.28 | ND |
| Macrophage | 0.44 | 0.58 | ND | ND | ND | ND | ND |
| T cell CD4+ | 0.41 | -0.44 | ND | ND | ND | ND | ND |
| T cell CD8+ | 0.16 | 0.48 | -0.27 | ND | -0.3 | 0.44 | ND |
| NK cell | -0.06 | ND | ND | ND | -0.06 | 0.42 | ND |
| Cancer associated fibroblast | 0.03 | ND | ND | ND | ND | -0.17 | 0.26 |
| Myeloid dendritic cell | 0.57 | 0.11 | -0.17 | 0.28 | ND | -0.31 | ND |
| Neutrophil | ND | 0.29 | 0.27 | ND | 0.51 | -0.3 | ND |
| NK cell activated | ND | ND | -0.44 | 0.29 | ND | ND | ND |
| T cell follicular helper | ND | ND | -0.39 | ND | ND | ND | ND |
| Macrophage M2 | ND | ND | 0.31 | 0.24 | -0.07 | ND | 0.19 |
| NK cell resting | ND | ND | 0.3 | -0.15 | ND | ND | ND |
| T cell CD4+ memory resting | ND | ND | 0.26 | 0.15 | ND | ND | ND |
| Mast cell activated | ND | ND | -0.25 | 0.49 | ND | ND | ND |
| Mast cell resting | ND | ND | 0.23 | ND | ND | ND | ND |
| Myeloid dendritic cell resting | ND | ND | ND | 0.44 | ND | ND | ND |
| Monocyte | ND | ND | ND | 0.41 | 0.31 | -0.29 | ND |
| Macrophage M0 | ND | ND | ND | -0.24 | ND | ND | ND |
| B cell plasma | ND | ND | ND | -0.16 | ND | ND | ND |
| Macrophage M1 | ND | ND | ND | ND | 0.34 | ND | ND |
| T cell CD4+ (non-regulatory) | ND | ND | ND | ND | -0.28 | ND | ND |
| uncharacterized cell | ND | ND | ND | ND | -0.21 | ND | ND |
| T cell regulatory (Tregs) | ND | ND | ND | ND | -0.2 | ND | ND |
| Macrophage/Monocyte | ND | ND | ND | ND | ND | -0.29 | ND |
| T cell | ND | ND | ND | ND | ND | 0.27 | ND |
| Hematopoietic stem cell | ND | ND | ND | ND | ND | ND | 0.32 |
| T cell CD4+ Th2 | ND | ND | ND | ND | ND | ND | -0.31 |
| Endothelial cell | ND | ND | ND | ND | ND | ND | 0.24 |
| T cell CD8+ naive | ND | ND | ND | ND | ND | ND | -0.22 |
| T cell CD4+ Th1 | ND | ND | ND | ND | ND | ND | -0.22 |
| Common lymphoid progenitor | ND | ND | ND | ND | ND | ND | -0.2 |
| Plasmacytoid dendritic cell | ND | ND | ND | ND | ND | ND | -0.19 |

**Table S2:** Comparison of clustering with and without immune inference. Left side: patients are clustered based on expression without immune inference. Right side: patients are clustered after immune inference. The number of clusters is dependent on the uniqueness of each method of estimation of immune inference.

| Immune Inference |  |  |  |  |  |
| --- | --- | --- | --- | --- | --- |
| Without |  |  | With |  |  |
| KRAS |  |  |  |  |  |
| Sub0 | Mutated | 63 | Sub0 | Mutated | 48 |
|  | WT | 146 |  | WT | 116 |
| Sub1 | Mutated | 8 | Sub1 | Mutated | 16 |
|  | WT | 40 |  | WT | 59 |
| Sub2 | Mutated | 57 | Sub2 | Mutated | 86 |
|  | WT | 94 |  | WT | 178 |
| Sub3 | Mutated | 22 |  |  |  |
|  | WT | 73 |  |  |  |
| EGFR |  |  |  |  |  |
| Sub0 | Mutated | 30 | Sub0 | Mutated | 33 |
|  | WT | 179 |  | WT | 131 |
| Sub1 | Mutated | 6 | Sub1 | Mutated | 4 |
|  | WT | 89 |  | WT | 71 |
| Sub2 | Mutated | 18 | Sub2 | Mutated | 25 |
|  | WT | 133 |  | WT | 239 |
| Sub3 | Mutated | 8 |  |  |  |
|  | WT | 40 |  |  |  |

## Disclaimer/Publisher’s Note

The statements, opinions and data contained in all publications are solely those of the individual author(s) and contributor(s) and not of MDPI and/or the editor(s). MDPI and/or the editor(s) disclaim responsibility for any injury to people or property resulting from any ideas, methods, instructions or products referred to in the content.

